# Bioclimatic variables influence the strength of purifying selection on mitochondrial DNA in an avian clade (Aves: Piciformes)

**DOI:** 10.64898/2026.06.11.731604

**Authors:** Jérôme Fuchs, Benoit Nabholz, Brieuc Kaesmann, Jean-Marc Pons, Céline Bonillo, Martin Irestedt, Sophea Chhin, Dawie de Swardt, Inocencio Chongo, Almiro Tivane, Eduardo Samo Gudo, Per G. P. Ericson

**Affiliations:** Institut de Systématique, Evolution, Biodiversité (ISYEB), Muséum national d’Histoire naturelle, CNRS, Sorbonne Université, EPHE, Université des Antilles, CP51, 57 rue Cuvier, 75005 Paris, France; ISEM, Univ Montpellier, CNRS, IRD, Montpellier, France; Institut Universitaire de France (IUF), Paris, France; Department of Bioinformatics and Genetics, Swedish Museum of Natural History, Stockholm, 114 18, Sweden; Department of Biodiversity, General Directorate of Policy and Strategy, Ministry of Environment, Cambodia; Department of Ornithology, National Museum, P. O. Box 266, Bloemfontein, South Africa; Instituto Nacional de Saúde, Mozambique

**Keywords:** Piciformes, mitogenome, molecular evolution, bioclimatic variables

## Abstract

Mitochondrial loci were for long considered as markers of choice to reconstruct phylogenies. The development of high-throughput sequencing over the past two decades fostered the sequencing of mitogenomes, allowing further macroevolutionary questions to be tested. Several biological traits of birds (e.g. body mass, migration distances) have been related to mitochondrial substitution rates. Environmental parameters in ectothermics vertebrates, and potentially in endotherms, have been further suggested to impact substitution rates for specific taxa or loci. Yet, the relative importance of biological traits versus bioclimatic variables is unknown because the former were not systematically controlled for in studies that underlined the effect of the bioclimatic variables. To assess the importance of bioclimatic variables on selection regimes, we analysed the thirteen mitochondrial protein-coding genes for 176 Piciformes (toucans, honeyguides, woodpeckers), a clade with homogeneous life-history traits that can be found in diverse bioclimatic environments. Our analyses highlighted a negative relationship between temperature annual range and the non synonymous to synonymous substitutions ratio. The higher purifying selection in temperate environments may be a result of the strong constraints on maintaining an optimal metabolism in broader climatic variations. Our results further highlight that care should be taken when applying ‘general’ mitochondrial clocks to estimate divergence times among avian lineages distributed in different climatic conditions.

## Introduction

The avian mitochondrial genome is a non-recombining, maternally inherited, circular molecule that consists of 37 genes, among which 13 code for proteins, as well as 22 transfert RNAs (tRNAs), 2 ribosomal RNAs (rDNAs) and one or two control regions. The thirteen proteins are present in four primary metabolic pathways (ND-, CO-, ATP and cytochrome b) from the Krebs cycle (Scheffler 2007). Mitochondrial DNA was for long considered as a marker of choice for phylogenetic and phylogeographic studies because of its ease to amplify, high variability both among and within species, and claimed neutrality of its selective regime (but see Ballard and Whitlock 2004, Galtier et al. 2009). Over the last two decades, several life history traits or biological characteristics of vertebrates have been significantly related to mitochondrial substitution rates or selective pressures: longevity (Nabholz et al. 2009), body mass (Nabholz et al. 2016), metabolic rate (Chong and Mueller 2012), migration distance (Pegan et al. 2024), life style (e.g. feeding habits in snakes Castoe et al. 2009, aquatic life in turtles, Escalona et al. 2017; blood feeding in bats, Botero-Castro et al. 2018), loss of flight in birds (Shen et al. 2009) or gain in bats (Shen et al. 2010), and environmental parameters in ectotherms (e.g. turtles, Lourenço *et al*. 2013) and potentially in endotherms (mammals, Gilman et al. 2009; birds: Bleiweiss 1998, Ribeiro et al. 2011, Ramos et al. 2018). It is nevertheless difficult to draw any definitive conclusion concerning the relative importance among all the above mentioned biological effects because the corresponding studies adopted different sampling strategies (closely or more distantly related species, number of species pairs) and/or methods (species pairs or whole clade, phylogenetic versus non-phylogenetic methods) (Orton et al. 2019).

The Piciformes (jacamars, puffbirds, barbets, toucans, honeyguides, woodpeckers, 449 species, www.worldbirdnames.org) is a monophyletic lineage of birds that diversified over the last 50 millions years in both temperate and tropical regions (e.g. Prum et al. 2015, Shakya et al. 2017); some members of this clade, the Picidae, are well known for their drumming as a signaling behaviour (e.g. Garcia et al. 2020). Piciformes are overall poor dispersers and only a very few Picidae species from temperate environments (e.g. *Jynx torquilla*, *Sphyrapicus* ssp, *Melanerpes erythrocephalus*) show migratory movements. Most Piciformes species have an omnivorous diet (carnivorous and frugivorous) during their annual cycle. The Piciformes thus consists of a rather homogenous group for some of the life history traits and biological characteristics (e.g. migration distance, flight ability, diet) that have been related to mitochondrial substitution rate variations in vertebrates.

In the present study, we used the Piciformes order as a model clade to aimed to 1) test the level of selection acting on mitochondrial protein-coding genes across the Piciformes and 2) determine whether two more variable ecological/biological characteristics (climatic variables, body mass) within the Piciformes influence the dN/dS ratio.

## Material and methods

### Sampling

Most of the Piciformes families are only distributed in a single major biogeographic area (e.g. Central/South America for the Bucconidae, Galbulidae, Ramphastidae; South-East Asia for the Megalaimidae). In contrast, the Picidae (239 species) have a much wider distribution, being present in all biogeographic areas but Australasia, Madagascar and the Oceanic islands. Consequently, the most speciose family the Picidae was more densely sampled (Supplementary Table S1, Prum et al. 2015, Tamashiro et al. 2018). Ultimately, the final data set consists of 176 species level taxa. The taxonomy followed the IOC World Bird list v.13.2 (www.worldbirdnames.org)

### DNA extraction

We extracted DNA from tissue (muscle, liver), blood using the Qiagen extraction kit (Qiagen, Valencia, CA) following the manufacturer’s protocol. For the toe pad samples, DNA was extracted and purified using either a Phenol-Chloroform protocol or a CTAB-based protocol (Winnepenninckx et al. 1993). DNA quality and purity for samples sequenced using next-generation sequencing were assessed using a Qubit Quantification (Qubit dsDNA BR Assay Kit, Life Technologies), NanoDrop spectrophotometer (NanoDrop Technologies, Inc., Wilmington, DE) and Fragment Analyzer (Agilent).

### Amplification, sequencing and assemblies

We used four strategies to obtain the mitochondrial genomes.

#### a) Amplicon sequencing

Overlapping 0.8-1.5 kb fragments were amplified from modern samples and Sanger-sequenced using a combination of universal and specific primers (Supplementary Table SX). This approach was complemented by performing overlapping long-range pcrs (range 4-8 kb) before being fragmented and sequenced either on an IonTorrent sequencer (Fuchs et al. 2016) or a Miseq Illumina sequencer using a mixed species pooling strategy (see Hinsinger et al. 2015). Assemblies were made using a mix strategy of de novo assemblies of *iontorrent* or *illumina* reads in Mitobim 1.9.1 (https://github.com/chrishah/MITObim; Hahn et al. 2013) preceded or not by read mapping and/or genome assemblies in *Ugene* Okonechnikov *et al*., 2012).

#### b) mtDna bait capture

We also obtained mitochondrial DNA sequence using a mitochondrial bait-capture approach. A custom bait-set (myBaits Target Capture Kit Cat# 300216, Ref# 190124–90, Daicel Arbor Biosciences, Ann Arbor, MI) was defined using a representative set of Piciformes lineages that were either published on Genbank or already obtained by us (cut-off date: 1 June 2017). The baits set included 14,081 baits (100 base pairs long). We used 500 ng of uniquely barcoded extracted DNA per sample as an input and pooled eight individuals per capture reaction. We followed the manufacturer’s protocol for the capture reactions excepted that we skipped the DNA fragmentation step since the historical DNA is already fragmented. Briefly, Hybridization reactions were prepared using the myBaits v.5.01 protocol, with 24 hours incubation at 63°C and used the NebNext ultra II Q5 master mix on-bead PCR method. We amplified the enriched libraries using 22 cycles of PCR and cleaned them using 1.15X SPRI beads. We determined target molarity and quality for sequencing using Qubit, and determined average fragment size using Bioanalyser High Sensitivity. Sequencing was performed on an Illumina MiSeq V2 sequencer at the ‘Institut du Cerveau et de la Moëlle Epinière’ (ICM, Pitié-Salpêtrière Hospital, Paris).

#### c) Whole genome shotgun sequencing

Ten samples (see Supplementary Table 1) were subject to genome shotgun sequencing from museum study skins (n=7) or modern samples (n=3). For the *Xiphidiopicus* historical sample, library preparation and indexing followed guidelines from Irestedt et al. (2022) and Meyer and Kircher (2010). The indexed library was sequenced on a half lane on the Illumina Hiseq X platform. Raw reads were cleaned by removing adapter contamination, low-quality bases and low complexity reads using a custom designed workflow (https://github.com/mozesblom) that utilizes TRIMMOMATIC v.0.32 (Bolger et al., 2014) using default settings. Cleaned reads were mapped using BWA mem (v.0.7.15; Li and Durbin, 2009) against the previously published complete mitochondrial genome of *Campephilus guatemalensis* (GenBank NC0028020). FreeBayes (v.1.1; Garrison and Marth, 2012) was used to call variants based on reads with a mapping quality above 10 and VCFfilter (https://github.com/vcflib) to exclude variant calls with a quality score below 20 or an allelic balance between 0 and 0.2. Moreover, VCFfilter was also used to create a mask file, which included sites with putative heterozygous positions or sites with a read depth coverage below 20X. Finally, combining both the variant and mask files, BCFtools (v.1.4.1; www.htslib.org) was used to call a consensus mitogenome. The six Megalaimidae toe pad samples that were subject to shotgun sequencing were sequenced on a NovaSeq 6000 SP sequencer at the ‘Institut du Cerveau et de la Moëlle Epinière’ (ICM, Pitié-Salpêtrière Hospital, Paris). We assembled the mitochondrial genomes for the six Megalaimidae species with NOVOPlasty-2.7.2 (Dierckxsens et al. 2017; genome range of 16500-19000 bp, K-mer of 39, insert size of 200 bp) and by mapping using the BWA algorithm, as implemented in Ugene (Okonechnikov *et al*., 2012). Mapping was performed using the closest *Psilopogon* species from the target species (den Tex and Leonard 2013).

#### d) Genbank sequences

Further mitochondrial genomes were obtained either directly from Genbank (www.ncbi.nlm.nih.gov), by de novo mitochondrial assemblies using NOVOPlasty-2.7.2 (Dierckxsens et al. 2017) or mapping using the BWA algorithm, as implemented in Ugene (Okonechnikov *et al*., 2012) from the raw short reads sequence data obtained from whole genome sequencing (n= 21; Manthey *et al*. 2018, Feng et al. 2020, Kimmitt et al. 2023, Pirro et al., unpublished) or as by-product of Ultra-Conserved Elements capture and sequencing (n=4; Bocalini et al. 2021, Smith et al. 2021; see Supplementary Table S1). In some cases, the published mitochondrial genomes missed ND6; we could assemble this locus for all species for which it was missing by using the corresponding Single Read Archives and the ND6 sequence of the closest species as a bait.

### Alignment and Phylogenetic analyses

Alignment of protein-coding genes were performed by eye using the amino acid translation and based on previously annotated mitochondrial genomes. All alignments used for the phylogenetic analyses are available at XXXXXXXXX. Newly generated sequenced have been deposited in Genbank (Accession numbers: XXXX-XXXX).

Divergence times on the concatenated data set were estimated using BEAST 1.10.4. (Suchard et al. 2018). We used the GTR+G+I substitution model for all loci (thirteen partitions), the substitution model selected in MEGA X (Kumar et al. 2018) under the BIC criterion, and assumed strict clock distributions. We specified a Birth-Death prior for the tree prior. MCMC chains were run for fifty million iterations with trees and parameters sampled every 5000 iterations.

The oldest well assigned fossils are *Piculoides saulcetensis*, from the early Miocene of France (−22.5 mya, De Pietri et al. 2011) and a feather from an unnamed taxon from the Dominican Republic that was preserved in amber (Laybourne et al. 1994). The former was suggested by the authors to either represent a stem Picidae or a stem Picinae + Picumninae clade. The latter fossil was considered to be more closely related to the Antillean Piculet than to any other Picidae (Laybourne et al. 1994). More fossils have been found from Pliocene or Pleistocene deposits. Most Pleistocene deposits refer to modern species so their use as calibration points for higher level phylogenies is difficult. The relationships of most Miocene/Pliocene fossils are uncertain (opt cit. Manegold and Louchart 2012). The only one exception being *Australopicus nelsonmandelai* from the Pliocene deposit of Langebaanweg in South Africa (Manegold and Louchart 2012); *Australopicus* has been suggested to be related to the genera *Dryocopus* and *Celeus* (Manegold and Louchard 2012), two genera that do not currently occur in sub-Saharan Africa. We decided to not use this calibration point because of 1) the lack of support for the relationships among the primary Picinae lineages and 2) the fact that not all primary lineages recovered in molecular analyses were retrieved in the morphological analyses of Manegold and Louchard (2012). We think that using *Australopicus* as a calibration point could be considered a very strong assumption because it would attract ‘basal’ divergence within the Picini toward the present. Instead, we used *Piculoides saulcetensis* as a calibration point. We used a lognormal Prior (mean=0.0, stdev=0.5 offset=22.5) for the split between the Jynginae and the remaining Picidae; this split implies that *Piculoides saulcetensis* is more likely situated at the base of the Picumninae/Picinae clade rather than at the base of the Picinae. Given the short internodes for the relationships among the three Picumninae lineages and the Picinae, we consider that the temporal bias in the choice of this calibration point is limited.

All phylogenetic analyses were run on the genotoul cluster (https://bioinfo.genotoul.fr/). We used TRACER v1.7 (Rambaut et al. 2014) to help ensure that the effective sample size for all Bayesian analyses of the underlying posterior distribution was adequate (>200) for meaningful estimation of parameters.

### Relationships between bioclimatic conditions and dN/dS

We used PAML 4.10.6 (Yang 1997, 2007) to test for positive or purifying selection acting on particular branches (branch model), sites (site model) and the combination of the two (branch-site models). All analyses were performed three times to ensure convergence in likelihood values. We particularly tested two hypotheses by flagging nodes or leaves on the PAML input tree. We tested whether bioclimatic factors have some impact on selection by coding each species as ‘temperate’ vs ‘tropical’, based on geography. The Principal Component Analysis performed on the 19 bioclimatic variables (Figure 1, see below) largely confirmed this distinction but also indicated that two species could be alternatively coded: *Dendrocopos noguchii* and *Veniliornis nigriceps* (Figure 1). We performed the analyses using the ‘geographic’ (hereafter referred to as ‘climate’) and the alternative coding (hereafter referred to as *climate alternate coding*) for these two species.

**Figure 1.**
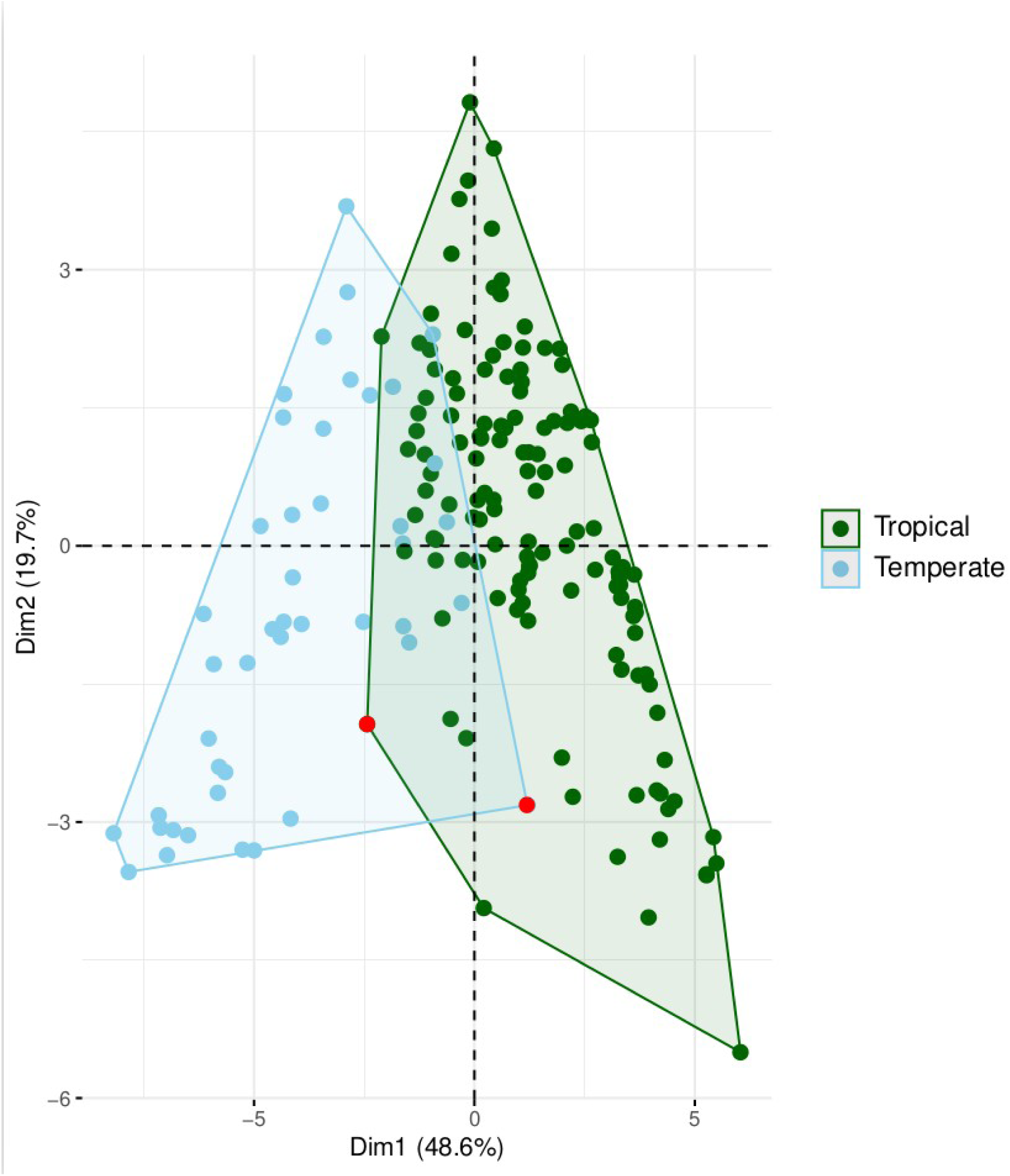
Projection of the principal components for the 163 species level taxa on the first two axes (51.8 % and 18.6 % of the total variation). Colours indicate species found in temperate (blue) or tropical (green) bioregions. The two species (*Dendrocopos noguchii* and *Veniliornis nigriceps*) with a strong mismatch between geography and climate are highlighted in red dots.

#### Effect of climate on dN/dS

The likelihood of a model with one dN/dS (ω) for the whole phylogeny was compared with a model with two dN/dS (one for the background branches and one for the branches with shifts in climatic category-temperate versus tropical). This test corresponds to the branch model. The analyses were done on each protein-coding gene individually as well as on the concatenated alignment.

#### Effect of climate on positive selection

We performed two nested model comparisons (M1a/M2a and M7/M8) that allows for a variable ratio of ω among sites across the entire phylogeny. Model M7 constrains ω to be between 0 and 1 for all sites, whereas M8 introduces an additional site class with ω > 1. Comparing the likelihoods of the M7 and M8 models enables the detection of positive selection in the alignment; this test does not require the specification of foreground branches and corresponds to the site model. The analyses were done on each protein-coding gene individually as well as on the concatenated alignment. We also compared the likelihood of a null model where no site is allowed to evolved with a ω > 1 (settings fix_omega = 1 and omega =1, model M1a) with an alternative model where a class of sites (proportion p2) are allowed to have a ω > 1 in particular ‘foreground’ branches (settings fix_omega = 0, model M2a) (Zhang et al. 2005); this test corresponds to the branch-site model of positive selection and requires the specification of foreground branches (here discrete climatic categories-temperate versus tropical). The analyses were done on each protein-coding gene individually as well as on the concatenated alignment.

### Correlation between continuous variables and substitution rates

We used *Coevol v1.6* (Lartillot and Poujol 2011) to assess 1) if there is any bioclimatic variable that could be more directly linked to the difference in dN/dS between temperate and tropical species and 2) if there is any effect body mass on dN/dS.

We extracted median values for the 19 bioclimatic variables listed on worldclim (https://www.worldclim.org/bioclim; 2.5° resolution) from species occurrence points using the packages *geodata 0.6-6* and *terra 1.8-93* (Hijmans 2026a, b). Species occurrence points were gathered using the *rgbif 3.8-4* package (Chamberlain et al. 2026) with the filters *hasCoordinate=TRUE* and *basisOfRecord = ‘preserved_specimen’*, as implemented in RStudio 2024.12.563 (Posit team 2025). Because global distribution data may contain some errors (e.g. inversion of latitude and longitude columns), we checked that occurrence data were consistent with known distribution of taxa; aberrant points (e.g. georeferencing of captive birds, inversion of latitude and longitude columns) were excluded for the bioclimatic data extraction step. Only one occurrence per georeferenced point was kept for bioclimatic data extraction step (using the *unique* function). For 12 species for which species level genetic differentiation was detected, we extracted distinct bioclimatic data for each species level lineages by using various filters. The only exception was *Dendropicos fuscescens hartlaubii* for which too few data were available; we used bioclimatic data extracted from localities belonging to this lineage based on phylogeographic level sampling for this species (J. Fuchs, unpublished). To reduce the number of continuous variables for the *Coevol analyses,* we performed a Principal Component Analysis (PCA) on the 19 bioclimatic variables using *FactoMiner* (Lê et al. 2008) and visualized the results of the PCA using *factoextra* (Kassambara and Mundt 2020) and *ggplot2* (Wickham 2016).

Body mass data were mostly obtained from the Elton Traits database (Wilman et al. 2014). In cases of recent splits (e.g. *Chrysocolaptes* ssp, *Campethera permista* from *C. maculosa*), we also made specific queries on vertnet (www.vertnet.org) or from our own field work (*D. analis*). Body mass data for *D. noguchii* was obtained from Kotaka (2011). When no mass data was available, we used the same mass as the species to which the splitted taxa was considered to belong.

The strength of the coupling between a molecular variable (dS, dN/dS) and continuous variables are measured by the correlation coefficient between the variables, and the statistical support is evaluated based on the posterior probability (pp) of a positive (pp close to 1) or negative covariation (pp close to 0). The ratio of non-synonymous to synonymous substitutions dN/dS and dS were analyzed as dependent variables. We ran two independent analyses of 5000 samples (burnin 1000 samples) and assessed convergence using the *tracecomp* function implemented in *Phylobayes* (Lartillot and Philippe 2004).

### Controlling for the effect of long term effective population size

Long term effective population size (Ne) has a strong effect on the efficiency of selection process. Indeed, purifying selection is more efficient in large population whereas genetic drift is more prevalent in population with smaller Ne. It is still currently unclear whether temperate or tropical species hold larger Ne. Higher latitude species have usually larger distributions and population sizes than tropical species, suggesting that Ne could be higher in temperate species. Yet, the effects of the Plio-Pleistocene climatic cycles on species range and population census were possibly more dramatic for temperate species than for tropical species suggesting that long term Ne could in fact be higher in the tropics.

We compared the effective population sizes of 25 Piciformes species for which genome-wide sequencing data and a reference genome from a conspecific or congeneric individual were available (Table S2) using the pairwise sequentially Markovian coalescent (PSMC, *psmc-0.6.5*) model (Li & Durbin 2011). We particularly aimed to test whether temperate or tropical have higher Ne. We used *Trimmomatic-0.39* (Bolger *et al*. 2014) to clean the reads downloaded from the European Nucleotide Archive (https://www.ebi.ac.uk/). Paired and unpaired reads were then mapped to their respective species reference assembly using *BWA-0.7.17* (Li & Durbin 2009); PCR duplicates were removed using *Picard-2.20.7* (Picard Toolkit 2019). We performed variant calling using *samtools-1.14* (mpileup) and *Bcftools-1.9* packages (Li *et al*. 2009) on the assembly and used the VCFutils.pl script to quality filter variants (using -d100 -D10). PSMC were run using the following options for the number of free atomic time intervals (-p): “2+2+25*2+4+6” or “2+2+25*2+4+6+10”. These parameters are either the by-default values, excepted for the first time window which was split into two windows (“2+2” instead of “4”, Hilgers *et al*. 2025), or represent the values used by Nadachowska-Brzyska *et al*. (2015) with the exception of the first time window (Hilgers *et al*. 2025). We tested two values (5 or 15) for t – upper limit of time to most recent common ancestor- and set N to 25 and r to 5 (corresponding to defaults values). The distribution of Time to Most Recent Common Ancestor (TMRCA) was scaled into years by generation time and average mutation rate. We used a generation time of 2 years (Brommer *et al*. 2004) years and a mutation rate of 4.6 × 10^−9^ substitution/site/generation (Smeds *et al*. 2016) for all taxa. We tested for a difference in long term effective population size between tropical and temperate species by performing a Wilcoxon rank sum test in R version 3.6.3 (R Core Team 2020) on the median values of the four estimates for most recent Ne estimate for each species.

## Results

### Sequences characteristics

Our data set consisted of 176 species level lineages, among which 13 are not currently considered as distinct species by the IOC. Yet, these species-level divergences were either already highlighted in previous studies or matched plumage variation or known biogeographic regions (Supplementary Table S3).

The extra-nucleotide ND3-174 was found in all Piciformes lineages to the exception of the Megalaimidae (genera *Caloramphus* and *Psilopogon*) and five *Melanerpes* species (*M. a. aurifrons*, *M. a. santacruzi*, *M. carolinus, M. flavifrons* and *M. pucherani*). The remaining species had an extra C (wide majority), T (*C. nubica*, *D. benghalense*, *D*. *pyrrhogaster*, *M. tristis, M. candidus, M. cactorum, M. formicivorus, R. validus, I. maculatus*), or G (*B. macrodactylus*). Several insertion-deletion of codon were detected in the protein-coding genes (Supplementary Table S4).

### Phylogenetic relationships and divergence times

The final alignment of the thirteen protein-coding genes (stop-codon excluded) for the species-level phylogenetic analyses was 11,394 bp. The topology resulting from the partitioned concatenated analyses (Figure 2), was well supported as all higher-level relationships (among genera, tribes or families) but two were supported by posterior probabilities greater than 0.95. The two exceptions which were not strongly supported are 1) the relationships of *Xiphidiopicus* with respect to the genera *Melanerpes* and *Sphyrapicus* (PP: 0.48) and 2) the inter-relationships of *Dendrocopos*, *Dendropicos*/*Dendrocoptes*/*Leiopicus*/*Chloropicus* and *Dryobates*/*Veniliornis*/*Leuconotopicus* clades (PP: 0.72) and 3 the monophyly of the Lybiidae (PP: 0.91).

**Figure 2.**
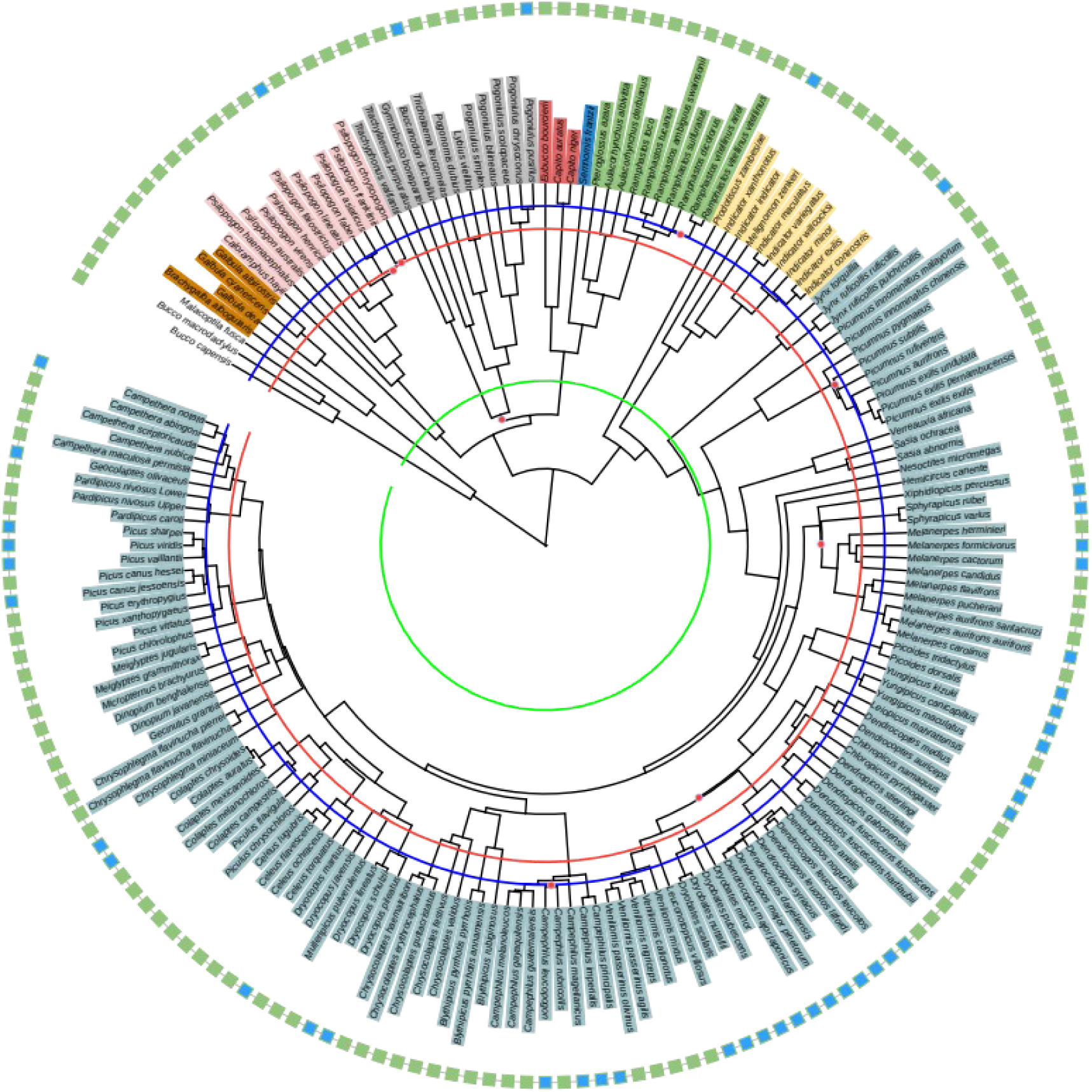
Maximum Clade Crediblity tree obtained using a partitioned concatenated scheme (176 individuals, 11,394 bp, 13 partitions). The fossil *Piculoides saulcetensis*, from the early Miocene of France (−22.5 mya, De Pietri et al. 2011) was used to calibrate the tree. Analyses were performed using BEAST 1.10.4. (Suchard et al. 2018) with a GTR+G+I substitution model for all loci (thirteen partitions), strict clock distributions for each loci and a Birth-Death prior for the tree prior. MCMC chains were run for fifty million iterations with trees and parameters sampled every 5000 iterations. Red circles indicate nodes supported by posterior probabilities smaller than 0.95.The Blue, Red and Green lines delimite the Pleistocene, Pliocene and Miocene epochs. Squares next no species names indicate the discrete climatic coding (green: tropical, blue: temperate).

The divergence time analyses (Supplementary Table S5) indicated that the Indicatoridae/Picidae diverged from the Megalaimidae/Capitonidae/Lybiidae/Semnornithidae/Ramphastidae (33.3 mya; 95% HPD: 31.5-35.1) at the same time than the Bucconidae diverged from the Galbulidae (32.0 mya; 95% HPD: 29.8-34.3), at the Eocene/Oligocene boundary. The Asian barbet Megalaimidae was the first lineage to split in the toucan/barbet clade, about 28.0 mya (95% HPD: 26.5-29.7). In the Indicatoridae/Picidae clade, the genus *Prodotiscus* was the first to branch off, about 27.8 mya (95% HPD: 26.3-29.3). The Picidae diverged from their closest relatives (*Indicator/Melignomon*) about 25.9 mya (95% HPD: 24.8-27.2). The wrynecks splitted from the remaining Picidae lineages about 23.5 mya (95% HPD: 22.8-24.4), followed by the sequential split of *Vivia*/*Picumnus* (20.7 mya, 95% HPD: 19.7-21.8), *Sasia*/*Verreauxia* (18.6 mya, 95% HPD: 17.6-19.7) and *Nesoctites* (14.7 mya, 95% HPD: 13.9-15.7). The Picinae radiated about 13.8 mya (95% HPD: 13.-14.6). The four primary Picinae lineages diversified shortly after about 9-11 mya (Campephilini, 10.9 mya, 95% HPD: 10.2-11.5; Picini 9.6 mya, 95% HPD: 9.0-10.1; Dendropicini 10.2 mya, 95% HPD: 9.6-10.8; Melanerpini 10.1 mya, 95% HPD: 9.4-10.6).

### Relationships between discrete bioclimatic conditions and dN/dS

#### Effect of the climate on dN/dS along the phylogeny

For species that live in temperate habitat, the dN/dS significantly decreased in ND2 (ω_0_=0.05147, ω_1_=0.04382, p=0.04; alternate coding: ω_0_=0.05143, ω_1_=0.04374, p=0.04, Table 1 and Supplementary Table S6) and for the complete alignment (ω_0_=0.04170, ω_1_=0.03741, p<0.0004; alternate coding: ω_0_=0.04171, ω_1_=0.03716, p<0.00002) but increased in ND3 (ω_0_=0.04315 ω_1_=0.06974, p<0.001; alternate coding: ω_0_=0.04328, ω_1_=0.06948, p<0.001) (Table 1 and Supplementary Table S6). Coding had some impact as a significant impact of the climate was detected for CO1 (ω_0_=0.00592, ω_1_=0.00433, p=0.04) and ND5 (ω_0_=0.04158, ω_1_=0.04770, p=0.02) but the sigificance disappeared using the alternative coding (CO1: ω_0_=0.00588, ω_1_=0.00454, p=0.10; ND5: ω_0_=0.04172, ω_1_=0.04690, p=0.05). Altogether, these results suggest that purifying selection on the mitochondrial protein-coding genes is overall stronger in temperate species than in tropical species.

**Table 1.** Effect of climatic regime (temperate versus tropical) on dN/dS using the alternate coding for *Veniliornis nigriceps* and *Dendrocopos noguchii*. Lines in bold refer to loci for which the climatic regime had a significant effect (p<0.05).

| Loci | LnL | $\omega$ | Ln | $\omega_0$ | $\omega_1$ | p |
| --- | --- | --- | --- | --- | --- | --- |
| **All loci** | **-492954.460121** | **0.04112** | **-492945.048549** | **0.04171** | **0.03716** | **<0.00002** |
| ATP6 | -30528.963875 | 0.02982 | -30528.333068 | 0.02931 | 0.03354 | 0.26 |
| ATP8 | -8757.287406 | 0.15340 | -8757.114685 | 0.15523 | 0.14045 | 0.56 |
| CO1 | -52438.297131 | 0.00572 | -52436.924290 | 0.00588 | 0.00454 | 0.10 |
| CO2 | -24947.121747 | 0.01873 | -24947.113617 | 0.01877 | 0.01840 | 0.90 |
| CO3 | -29447.275584 | 0.02085 | -29446.492053 | 0.02127 | 0.01810 | 0.21 |
| Cytochrome b | -46145.299577 | 0.01993 | -46145.268257 | 0.02000 | 0.01951 | 0.80 |
| ND1 | -40425.985507 | 0.02215 | -40425.985240 | 0.02214 | 0.02219 | 0.98 |
| **ND2** | **-50819.783135** | **0.05034** | **-50817.625445** | **0.51430** | **0.04374** | **0.04** |
| **ND3** | **-15231.118744** | **0.04616** | **-15225.696839** | **0.04328** | **0.06948** | **<0.001** |
| ND4 | -63386.712570 | 0.03342 | -63386.712536 | 0.03342 | 0.03440 | 0.99 |
| ND4L | -13070.165371 | 0.03490 | -13069.785631 | 0.03423 | 0.03938 | 0.38 |
| ND5 | -85795.088154 | 0.04241 | -85793.221661 | 0.04172 | 0.04690 | 0.05 |
| ND6 | -24185.412225 | 0.05868 | -24185.391618 | 0.05849 | 0.05998 | 0.84 |

#### Effect of climate on positive selection

None the M1a-M2a, comparisons were significant. In contrast, the M7-M8 comparisons were significant for the concatenated alignment (p<0.0001), as well as for some individual loci (ND4: p<0.00001, ND6: p<0.00001; Supplementary Table S7). Thus, the PAML analyses indicate that positive selection may occur in the sequences but there is no evidence that it is related to bioclimatic variables.

### Correlation between substitution rates and continuous bioclimatic variables

The correlation between substitution rates (dS and dN/dS) and continuous variables were performed using five continuous variables: body mass and four bioclimatic variables retained after examination of the results of the Principal Component Analysis. The four bioclimatic variables were BIO7 (Temperature Annual Range), BIO8 (Mean Temperature of Wettest Quarter), BIO15 (Precipitation Seasonality -Coefficient of Variation-) and BIO17 (Precipitation of Driest Quarter). The first PCA Axis, mostly reflective of temperature seasonality, accounted for 48.6% of the variation, whereas the second PCA axis, mostly reflecting annual precipitation, accounted for 19.7%.

dS was negatively correlated with the dN/dS (pp=0, marginal correlation coefficient=-0.494) and with BIO8 (pp=0, marginal correlation coefficient=-0.722). The dN/dS was negatively correlated with BIO7 (Temperature Annual Range) (pp=0.005, marginal correlation coefficient=-0.382) (Table 2). Further posterior probabilities reached the significance threshold when analysing the partial correlations using the precision matrix; the two most supported correlations were the negative relationship between dS and BIO8 (pp=0, partial correlation coefficient=-0.835) and the negative relationship between dN/dS and BIO7 (pp=0, partial correlation coefficient=-0.799) (Table 2). These results suggest that species living in areas where annual temperature range is higher encompass stronger purifying selection.

**Table 2.**
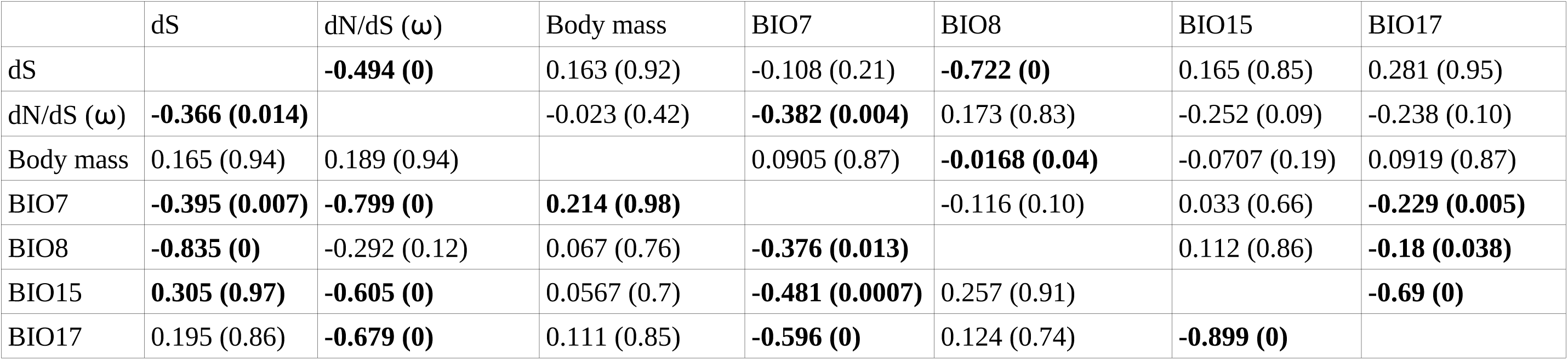
Results of the *Coevol* analyses (above diagonal -marginal correlation coefficients and associated p-values; below diagonal -partial correlation coefficients obtained using the precision matrix and associated p-values). Significant relationships (pp>0.95 or <0.05) are indicated in bold. Acronyms/parameters are: dS synonymous rate; dN/dS: non-synonymous to synonymous mutation rate, Body mass: average mass of individuals; BIO7 (Temperature Annual Range), BIO8 (Mean Temperature of Wettest Quarter), BIO15 (Precipitation Seasonality -Coefficient of Variation-) and BIO17 (Precipitation of Driest Quarter).

| | dS | dN/dS ($\omega$) | Body mass | BIO7 | BIO8 | BIO15 | BIO17 |
| --- | --- | --- | --- | --- | --- | --- | --- |
| dS |  | **-0.494 (0)** | 0.163 (0.92) | -0.108 (0.21) | **-0.722 (0)** | 0.165 (0.85) | 0.281 (0.95) |
| dN/dS ($\omega$) | **-0.366 (0.014)** | | -0.023 (0.42) | **-0.382 (0.004)** | 0.173 (0.83) | -0.252 (0.09) | -0.238 (0.10) |
| Body mass | 0.165 (0.94) | 0.189 (0.94) |  | 0.0905 (0.87) | **-0.0168 (0.04)** | -0.0707 (0.19) | 0.0919 (0.87) |
| BIO7 | **-0.395 (0.007)** | **-0.799 (0)** | **0.214 (0.98)** |  | -0.116 (0.10) | 0.033 (0.66) | **-0.229 (0.005)** |
| BIO8 | **-0.835 (0)** | -0.292 (0.12) | 0.067 (0.76) | **-0.376 (0.013)** |  | 0.112 (0.86) | **-0.18 (0.038)** |
| BIO15 | **0.305 (0.97)** | **-0.605 (0)** | 0.0567 (0.7) | **-0.481 (0.0007)** | 0.257 (0.91) |  | **-0.69 (0)** |
| BIO17 | 0.195 (0.86) | **-0.679 (0)** | 0.111 (0.85) | **-0.596 (0)** | 0.124 (0.74) | **-0.899 (0)** |  |

Body mass was negatively correlated with BIO8 (pp=0.04, marginal correlation coefficient=-0.165) but not with the dN/dS (Table 2); This relationship was not significant anymore when analysing the partial correlations (pp=0.76, partial correlation coefficient=0.067). Instead, body mass was correlated with BIO7 (pp=0.98, partial correlation coefficient=0.214). This result indicates that species living in areas where temperature is more variable (higher latitudes) are heavier, consistent with Bergmann’s rule.

### Controlling for the effect of long term effective population size

Our analyses detected a climate (temperate or tropical) effect on long term effective population size for the 25 analysed Piciformes (Supplementary Table S2 ; Wilcoxon rank-sum test: W = 39, p-value = 0.04). Excluding the Okinawa Woodpecker, because of its restricted and atypical distribution range for a Piciformes species, reinforced this trend (Supplementary Table S2; Wilcoxon rank-sum test: W = 29, p-value = 0.02).

Altogether, these results suggest that: 1) body mass is not related to the dN/dS in Piciformes and 2) species that are distributed in areas where temperature annual range is high encompass stronger purifying selection on mitochondrial protein-coding genes, 3) there is no evidence of positive selection linked to bioclimatic variables, and 4) tropical species have a tendency to have larger long term effective population size.

## Discussion

Mitochondrial DNA was for often considered as a neutrally evolving molecule (but see Ballard and Whitlock 2004, Galtier et al. 2009), making it a marker of choice to reconstruct phylogenies. Selection on the mitochondrial protein-coding genes has been a major research topic over the last twenty years (James et al. 2015) and since then have several life history traits or biological characteristics of vertebrates (Castoe et al. 2009, Nabholz et al. 2009, Shen et al. 2009, Shen et al. 2010, Chong and Mueller 2012, Nabholz et al. 2016, Escalona et al. 2017; Botero-Castro et al. 2018) and environmental parameters (Gilman et al. 2009, Lourenço *et al*. 2013, Ribeiro et al. 2011, Ramos et al. 2018) been suggested to be related to mitochondrial substitution rates. In the present study, we investigated the role of climate in shaping the selection regimes acting on the mitochondrial protein-coding genes in the avian order Piciformes.

### Climactic influence on mitochondrial DNA evolution

Our results suggest that climate plays a role in the mitochondrial DNA evolution with temperate species being under significantly stronger purifying selection than tropical species. The PAML analyses highlighted lower dN/dS (stronger purifying selection) in temperate species for the concatenated data whereas the COEVOL analyses revealed a significant negative relationship between dN/dS and annual temperature range.

Life history traits, such as active flight abilities could be a co-variate of climatic variables and annual temperature variation in particular. During the colder times of the year, food resources may be scarcer leading individuals to disperse via active flight to distant wintering areas. This migratory phenomenon, largely present in birds, is a highly energy consuming activity and possibly influence mitochondrial DNA evolution (Pegan et al. 2024). On the contrary, the evolution of flightlessness in some Galliformes resulted in the relaxation of selective constraints in mitochondrial protein-coding-genes (Shen et al. 2009). Very few woodpecker species from temperate environments (e.g. *Jynx torquilla*, *Sphyrapicus* ssp, *Melanerpes erythrocephalus*) have migratory movements. Hence, differences in flying capabilities or migratory behaviour may not explain the difference in the strength of purifying selection we observe in the Piciformes.

Another factor that could have resulted in a differential substitution dynamics of mitochondrial DNA between the temperate and tropical species is the differences in effective population size (Ne). For example, in case of a low long term effective population size, an increase of the dN/dS is expected due to a stronger effect of genetic drift in smaller populations (Ohta 1992, Woolfit 2009, Leroy et al. 2021). The long term effective population size of species occurring in temperate regions may have been more strongly impacted by the Plio-Pleistocene climatic cycles, through shrinkage of the extent of suitable habitat during glacial periods, than the effective population size of species in tropical regions where population extinction rates, and population size variations may have been lower, resulting in higher long term Ne in the tropics. On the opposite, lower latitude species could have smaller Ne, despite more stable long term effective population size, due to smaller distribution ranges. Empirical evidences for these theoretical assumptions are limited. For example, the long term Ne of one of the most numerous bird species in the world, the Palearctic European Starling (*Sturnus vulgaris*) is lower than the long term Ne of the eastern African endemic Superb Starling (*Lamprotornis superbus*) (Fuchs et al. 2024). For Piciformes, we detected a significant effect of latitude on long term with the Ne of tropical species being larger than the Ne of temperate species, implying that purifying selection should be more efficient in tropical species. Our analyses revealed that purifying selection is stronger in temperate species. Pegan et al. (2025) highlighted that temperate migratory species have larger long term effective population size than resident temperate species. Hence, for migratory species, it is possible that life history traits are more important than latitude in explaining differences in long term effective population size.

The mitochondrial genome codes for proteins that are involved in energetic production through the oxidative phosphorylation (OXPHOS) system. Selection on mitochondrial loci linked to climatic variables has already been highlighted within various bird (e.g. Ribeiro et al. 2011, Scott et al. 2011, Pavlova et al. 2013, Morales et al. 2015, Lamb et al. 2018, but see Orton et al. 2019) and mammal species (Gilman et al. 2009, Chen et al. 2018). In line with present study, Chen et al. (2018) found significant differences in the extent of purifying selection among temperate and tropical populations of the Wild Boar (*Sus scrofa*), with the Siberian populations having a significantly lower dN/dS than the Vietnamese population. Stager et al. (2014) also found a significant effect of latitude in the mitochondrial genome evolution of *Tachycineta* swallows (Hirundinidae) and concluded that the pace of life (shortest lifespan, higher reproductive rate in temperate species) could impact selective regimes. Yet, *Tachycineta* swallows occurring in temperate climate are also migratory and given the impact of the migratory behaviour on selective pressures (Toews et al. 2013, Pegan et al. 2024), it is unclear whether the differences in selective regimes between temperate and tropical species are better explained by pace of life, as suggested by Stager et al. (2014), or migratory behaviour.

The significant differences in strength of purifying selection among Piciformes distributed in temperate or tropical climate may either be linked to a stronger constraint in maintaining an optimal metabolism in broader climatic variations in temperate environments (see also Ribeiro et al. 2018 about thermosensation flexibility) and/or to accelerated evolution in a tropical environment. Gilman et al. (2009) found that substitution rates in the cytochrome b have been substantially faster for mammal species living in warmer habitats relative to species living in cooler habitats. As in the present study, these authors were able to exclude body mass or genetic drift as confounding factors. They proposed the ‘Red Queen Hypothesis’, which suggests that the rate of evolution of a species within a community dependent on the rate of evolution of other co-occurring species with which it is co-evolving (Gilman et al. 2009), in accordance with the ‘evolutionary speed hypothesis’ (Rohde 1992). Such hypothesis would imply that positive selection would be more common in tropical species than in temperate species. Evidences of positive selection in our data set were ambiguous. A few comparisons between the M7 and the M8 models were significant. Yet, none of these significant comparisons could be related to the bioclimatic variables as none of the M1a-M2a comparisons were significant. The M1a-M2a comparison is nevertheless more conservative than the M7-M8 comparison (Zhang et al. 2005). Consequently, it is currently difficult to conclude whether the significant signal for positive selection detected in the M7-M8 comparisons is the result of positive selection unrelated to bioclimatic variables or due to its less conservative nature.

Altogether our analyses suggest that the significant differences in strength of purifying selection among Piciformes distributed in temperate or tropical climate is better explained by stronger constraints in maintaining an optimal metabolism in broader climatic variations in temperate environments. Analyses of more clades distributed in temperate and tropical regions are needed to assess the generalisation of our result.

## Acknowledgments

We are very grateful to the following institutions and people for their invaluable contributions to our study: American Museum of Natural History, New York (J. Cracraft, P. Sweet) Field Museum of Natural History, Chicago (J. Bates, S. Hackett, D. Willard, B. Marks), Musée d’Histoire Naturelle, Genève (A. Cibois); Museum of Vertebrate Zoology, University of California, Berkeley (R.C.K. Bowie, C. Cicero); Louisiana State University, Museum of Natural Science, Baton Rouge (R. Brumfield, D. Dittmann, F.H. Sheldon); National Museum of Natural History, Washington (J. Dean, G. Graves); University of Washington, Burke Museum, Seattle (S. Birks, R. Faucett, J. Klicka), Zoological Museum, University of Copenhagen (J. Fjeldså, J.B. Kristensen). Laboratory work was performed at the UAR-2700 2AD (Service de Systématique Moléculaire). For help in the laboratory, we thank R. Debruyne, C. Ferreira, D. Gey, P. Sundararaju and J. Vasseur. We are grateful to the provincial authorities in the Eastern Cape, Limpopo, Kwazulu-Natal, Free State of South Africa, and Eastern Cape Parks for granting permission to collect samples and specimens (permits 0112-CPM401-00001, CPM-002-00003, OP 3771/2009, 01-24158, CRO144/14CR, FAUNA1066-2008, RA-0190) and the FitzPatrick Institute of African Ornithology (University of Cape Town) for help with logistics. We are also grateful the Limpopo Department of Economic Development, Tourism and Environmental Affairs (J. Heymans, T. J. Seakamela). We are very grateful to T. Saitoh (Yamashina Institute for Ornithology) for help with finding body mass data for the Okinawa Woopecker. Finally, we wish to thank the ATM “Biodiversité Actuelle et Fossile” and “Emergence” committees, the ‘Département Origines et Evolution’ and the ‘Société des Amis du Muséum’ for having funded this project and the National Museum of Maputo (Mozambique) for export permits.

## Ethics

Field sampling protocols were approved by Committte Cuvier d’éthique en matière d’expérimentation animale (DAECC 68-055 and 68-119).

## Author’s contributions

J.F.: conceptualization, data curation, formal analysis, investigation, methodology, validation, visualization, funding acquisition and writing—original draft; B.N.: conceptualization, methodology, formal analysis, validation, writing—review and editing; B.K.: formal analysis, writing—review and editing; J.M.P: writing—review and editing; C.B.: data acquisition, writing—review and editing; M.I.: data acquisition, writing—review and editing; S.C.: resources, writing—review and editing; D.d.S.: resources, writing—review and editing; I.C.: resources, writing—review and editing; A.T.: resources, writing—review and editing; E.S.G.: resources, funding acquisition, writing—review and editing; P.G.P.E.: funding acquisition, writing—review and editing.

All authors gave final approval for publication and agreed to be held accountable for the work performed therein.

## Conflic of Interest

We declare we have no competing interests.

## Data availability

All newly generated data have been submitted to Genbank (www.ncbi.nlm.nih.gov; Accession numbers XXXXX-XXXXX). The zenodo following DOI (10.5281/zenodo.20644470) includes 1) R scripts used to extract bioclimatic variables from occurrence data and generate Figure 1, 2) Input xml file used to generate Figure 2 and Maximum Clade Credibility tree in nexus format, 3) scripts used to generate population size history using the Pairwise Sequentially Markovian Coalescent (PSMC) model, 4) input and output files for the PAML analyses, and 5) input and output file for the coevol analyses

## SUPLPLEMENTARY MATEIRAL

### Material and Methods

#### Problems in published sequences

We detected several assembly or sequencing errors in sequences deposited in Genbank: 1) *Galbula dea* (MN356220) has a two base pair deletion in ND6 (positions 41-42), and 2) *Yungipicus kizuki* (OK011780) has a one base pair deletion in ND3 (position 137). These errors were manually corrected by deleting the one base insertion or by replacing gaps with Ns.

Finally, sequence inspections and comparisons with previously published data indicated that one of our previously published ND2 sequence (*Dryobates minor*, DQ188180, Fuchs et al. 2007) may be a numt as the newly sequenced we produced differs by 8 % from the previous sequence. Since the two sequences originate from different individuals sampled in Sweden (DQ188180) and France (this study), phylogeographic structure in this species could be an explanation. Yet, further mitochondrial loci for which sequences are available for both samples (e.g. CO1, ATP6, J. Fuchs unpublished) are much less divergent (CO1: 0.2%, ATP6: 0.3%). A Maximum Likelihood reanalysis of the ND2 data set in Fuchs et al. (2007) indicate that the *D. minor* sequences form a clade, suggesting that the numt is possibly specific to *Dryobates minor*.

#### Sequence characteristics of the analysed data set

When present, the amount of missing data varied between 1 bp (*Yungipicus kizuki*, 0.01%) and 623 bp (*Chrysocolaptes festivus*, 5.4%). Only three other species had more than 1% of missing data (*Celeus torquatus*, 377 bp, 3.3%; *Chrysocolaptes erythrocephalus*, 159 bp, 1.4%; *Trachyphonus purpuratus*: 196 bp, 1.7%).

Insertion or deletion of codon in protein (indels) occurred in eight out of the thirteen protein-coding loci (Supplementary Table S3). Some of the patterns were obviously different, as they imply a different number of codons, but appear to cluster in specific regions. Five out of the six cases in ND1 occur in the first 10 codons of the alignment, five out of the ten cases in ND4 occurred in the region 184 to 193 of the codon alignment and six out of the twelve cases in ND5 occurred in the region 23 to 37 of the codon alignment. Overall, indels occurred most frequently in the first or last positions of the codon alignment. Several indels were inferred to be homoplasic, usually shared between two unrelated lineages (for example *Celeus* and *Campephilus guatemalensis* share a three codon insertion in ND5).

**Supplementary Table S1.** List of samples analysed. Taxonomy follows IOC (2023). Asterisks indicate samples that were sequenced as part of this study. ** Specimens voucher in National Museum of Maputo (Mozambique). Acronyms are: AMNH, American Museum of Natural History, New York, U.S.A.; FMNH, Field Museum of Natural History, Chicago, U.S.A.; KU, University of Kansas Biodiversity Institute and Museum of Natural History, Lawrence, U.S.A.; LSUMNS, Louisiana State University, Museum of Natural Science, Baton Rouge, U.S.A.; MHNG, Musée d’Histoire Naturelle, Genève, Switzerland; MNCN, Museo Nacional de Ciencas Naturales, Madrid, Spain; MNHN, Muséum national d’Histoire Naturelle, Paris, France; MSB, Museum of Southwestern Biology, University of New Mexico, Albuquerque, U.S.A.; MVZ, Museum of Vertebrate Zoology, University of California, Berkeley, U.S.A.; NHMO, Natural History Museum, Oslo, Norway; NRM, Museum of Natural History Stockholm, Sweden; USNM, National Museum of Natural History, Smithsonian Institution, Washington, U.S.A.; UWBM, University of Washington, Burke Museum, Seattle, U.S.A.; ZMUC; Zoological Museum, University of Copenhagen, Copenhagen, Denmark. Individuals in bolld were sequenced as part of this study.

**Supplementary Table S2.**
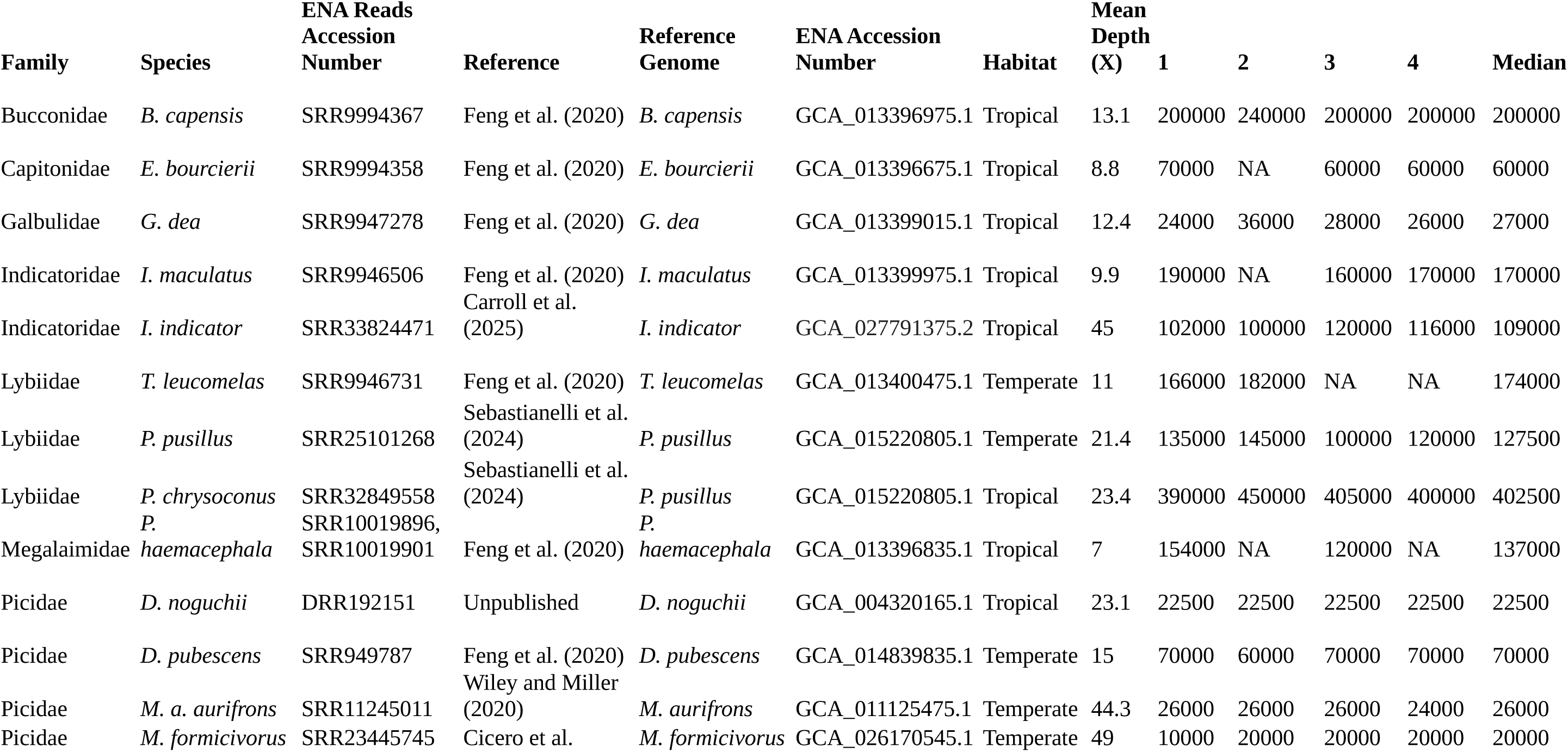
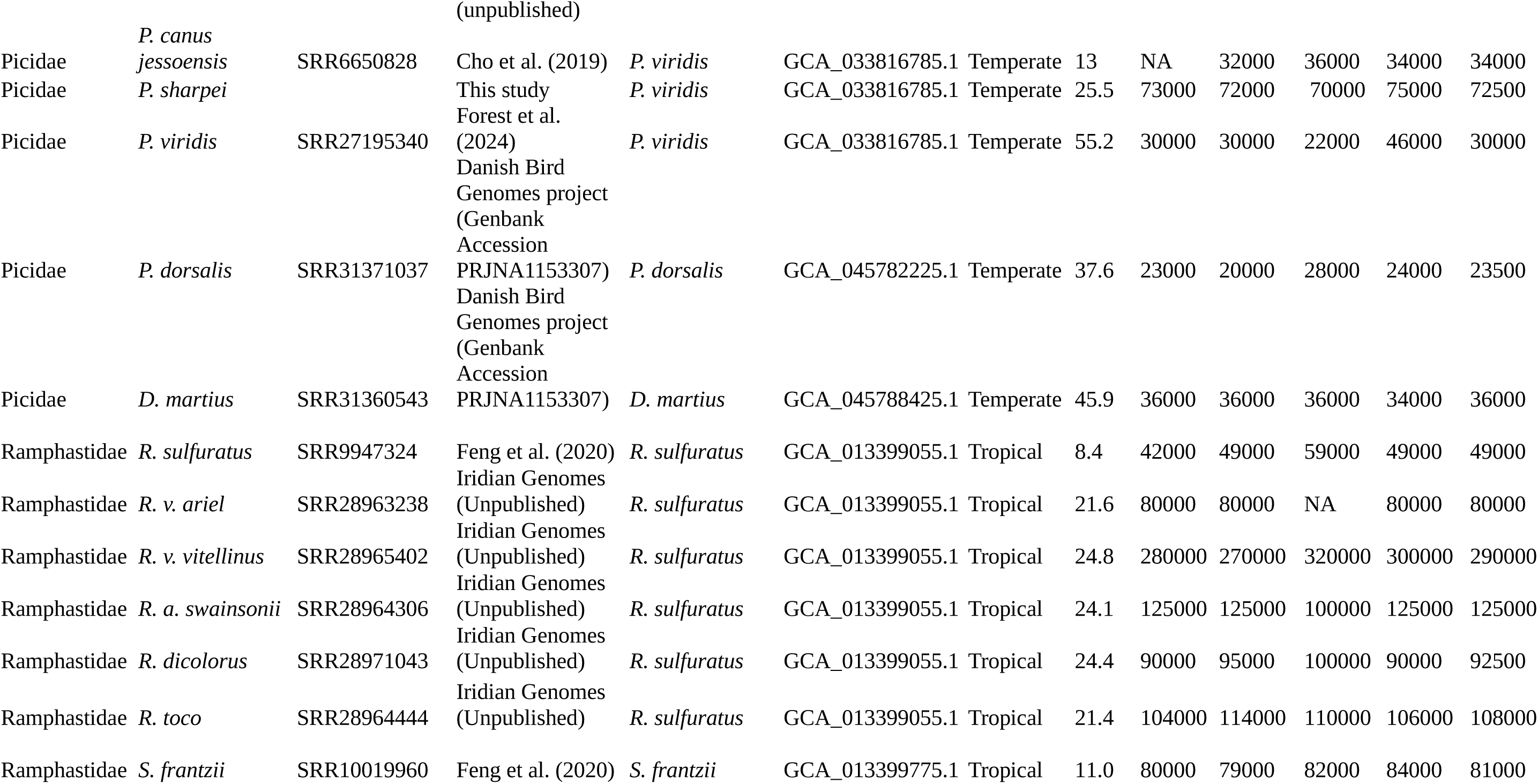
List of species included in the demographic analyses using the pairwise sequentially Markovian coalescent (PSMC, *psmc-0.6.5*) model (Li & Durbin 2011).. The demographic analyses were performed under the pairwise sequentially Markovian coalescent model. In all but two cases, reads were mapped on the reference genome from the same individual for which the reference genome was assembled. Ne values represent the most recent estimates. A Wilcoxon rank-sum test detected difference in Ne between temperate and tropical species (W = 39, p-value = 0.04). Parameters values for the PSMC analyses 1) -N25 -t5 -r5 -p “2+2+25*2+4+6”, 2) - N25 -t5 -r5 -p “2+2+25*2+4+6+10”, 3) -N25 -t15 -r5 -p “2+2+25*2+4+6”, 4) -N25 -t15 -r5 -p “2+2+25*2+4+6+10”. NA represent cases for which the estimates were highly divergent, and deemed dubious, from the three other estimates and caused by a very steep increase or decrease in Ne in the most recent times.

**Supplementary Table S3.** List of species within which substantial genetic divergence was high enough to consider them as distinct species-level lineages for the selection analyses.

| Species | Number of lineages | Divergence suggested before (taxonomy or molecular study) | Reference |
| --- | --- | --- | --- |
| *Blythipicus pyrrhotis* | 2 | Taxonomy |  |
| *Chrysophlegma flavinucha* | 2 | Taxonomy |  |
| *Dendrocopos leucotos* | 2 | Taxonomy/Molecular | Pons et al. (2021) |
| *Dendrocopos major* | 2 | Taxonomy/Molecular | Zink et al. (2002), Perktas et al. (2012) |
| *Dendropicos fuscescens* | 2 | Taxonomy/Molecular | Fuchs et al. (2017) |
| *Jynx ruficollis* | 2 | Taxonomy |  |
| *Pardipicus nivosus* | 2 | Taxonomy/Molecular | Fuchs and Bowie (2015) |
| *Picumnus exilis* | 3 | Taxonomy/molecular | Bocolini et al. (2020) |
| *Picumnus innominatus* | 2 | Taxonomy/molecular | Fuchs et al. (2007) |
| *Picus canus* | 2 | Taxonomy |  |
| *Veniliornis passerinus* | 2 | Taxonomy/Molecular | Shakya et al. (2017) |

**Supplementary Table S4.**
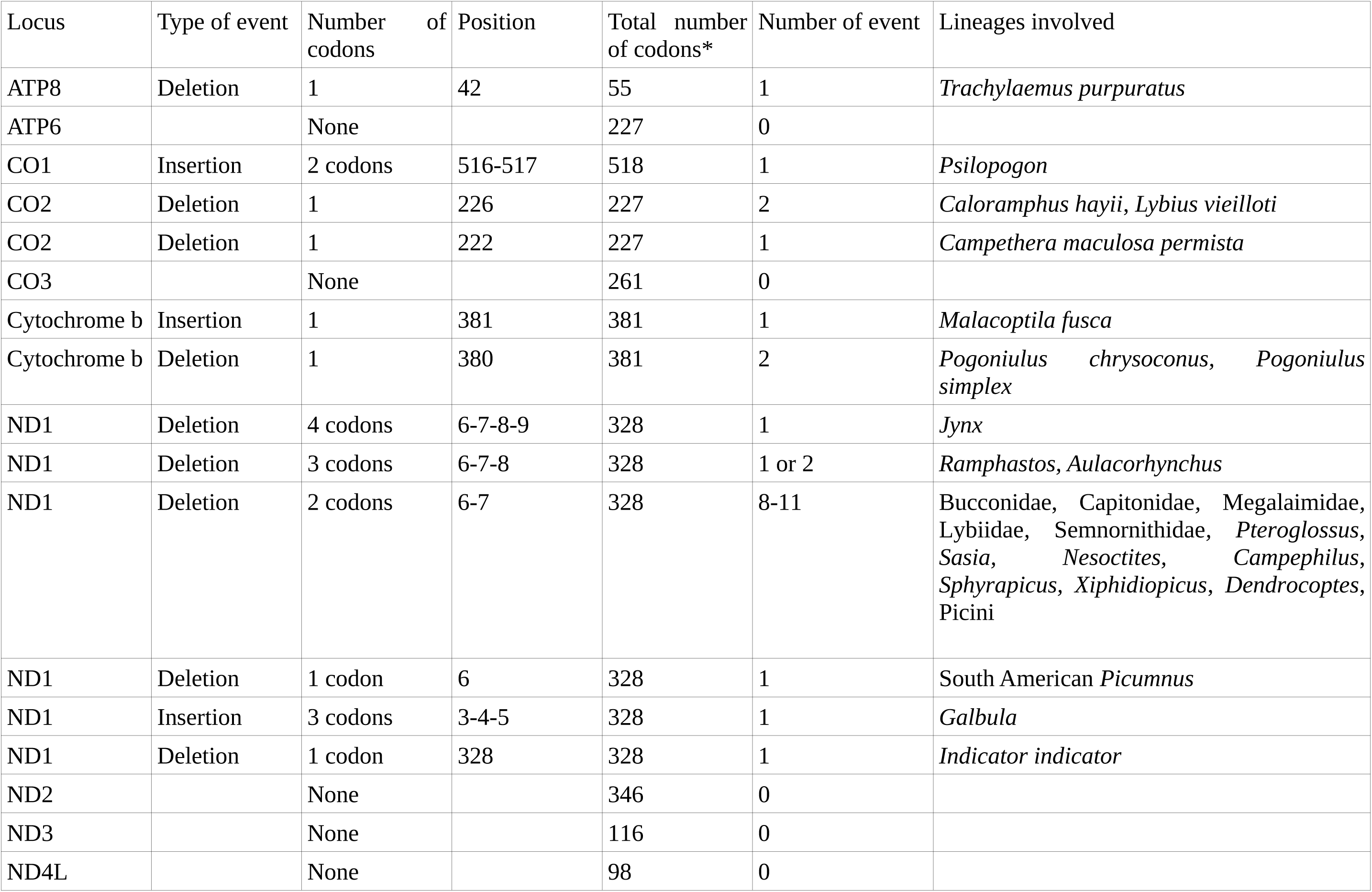
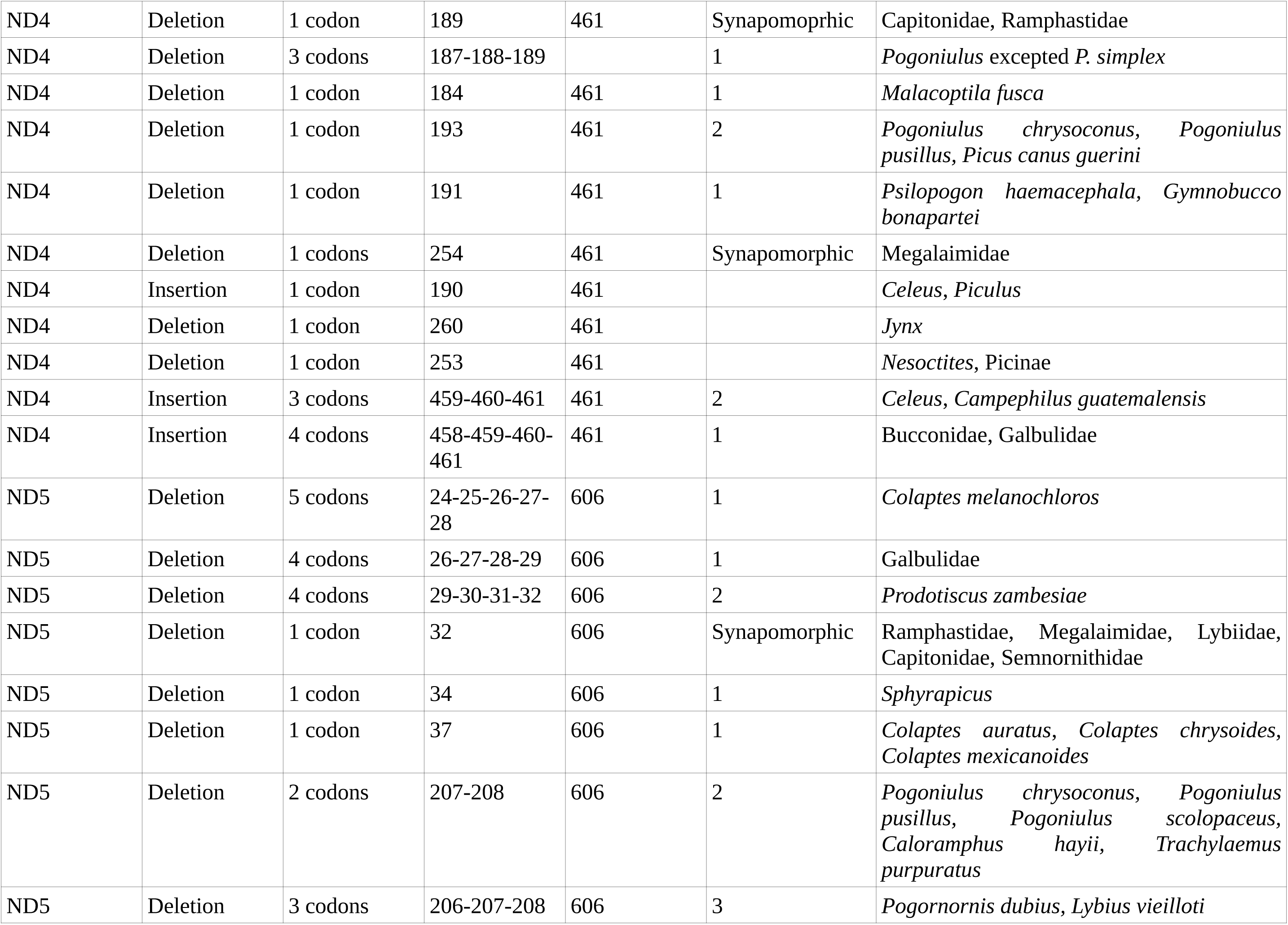
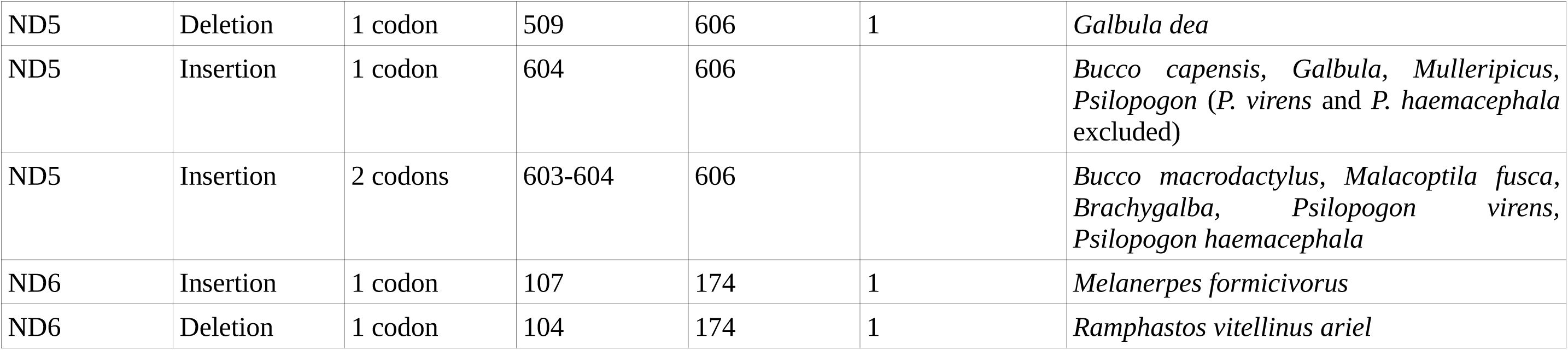
List of insertion deletion event in the protein coding involving a codon. * Excluding stop codon.

**Supplementary Table S5.** Substitution rates estimates for the Piciformes (this study) and, for comparison, with the estimates for a clade of Passeriformes (Lerner et al. 2011).

| Locus | Substitution rate – subs/site/myr/lineage-<br>(95% HPD) | Substitution rate – subs/site/myr/lineage-<br>(95% HPD) Lerner et al. (2011) |
| --- | --- | --- |
| ATP6 | 0.017 (0.015-0.0185) | 0.026 (0.021-0.031) |
| ATP8 | 0.017 (0.015-0.020) | 0.019 (0.013-0.024) |
| ND1 | 0.013 (0.012-0.0145) | 0.025 (0.022-0.029) |
| ND2 | 0.018 (0.016-0.02) | 0.029 (0.024-0.033) |
| ND3 | 0.017 (0.014-0.020) | 0.024 (0.018-0.031) |
| ND4 | 0.017 (0.016-0.019) | 0.022 (0.019-0.025) |
| ND4L | 0.0235 (0.020-0.027) | 0.025 (0.018-0.031) |
| ND5 | 0.0155 (0.0145-0.0166) | 0.021 (0.018-0.023) |
| ND6 | 0.017 (0.015-0.0195) | 0.024 (0.020-0.028) |
| CO1 | 0.015 (0.013-0.016) | 0.016 (0.014-0.019) |
| CO2 | 0.016 (0.014-0.0175) | 0.019 (0.016-0.022) |
| CO3 | 0.0145 (0.013-0.016) | 0.019 (0.015-0.021) |
| Cytochrome b | 0.0168 (0.015-0.0185) | 0.014 (0.012-0.016) |

**Supplementary Table S6.** Effect of climatic regime (temperate versus tropical) on dN/dS. Lines in bold refer to loci for which the climatic regime had a significant effect.

| Loci | LnL | $\omega$ | Ln | $\omega_0$ | $\omega_1$ | p |
| --- | --- | --- | --- | --- | --- | --- |
| **All loci** | **-492954.460121** | **0.04112** | **-492945.893303** | **0.04170** | **0.03741** | **<0.0004** |
| ATP6 | -30528.963875 | 0.02982 | -30528.327407 | 0.02930 | 0.03346 | 0.26 |
| ATP8 | -8757.287406 | 0.15340 | -8757.274242 | 0.15391 | 0.14987 | 0.87 |
| **CO1** | **-52438.297131** | **0.00572** | **-52436.239857** | **0.00592** | **0.00433** | **0.04** |
| CO2 | -24947.121747 | 0.01873 | -24947.119573 | 0.01875 | 0.01856 | 0.94 |
| CO3 | -29447.275584 | 0.02085 | -29446.472379 | 0.02128 | 0.01811 | 0.21 |
| Cytochrome b | -46145.299577 | 0.01993 | -46145.288598 | 0.01997 | 0.01969 | 0.88 |
| ND1 | -40425.985507 | 0.02215 | -40425.941281 | 0.02205 | 0.02274 | 0.77 |
| **ND2** | **-50819.783135** | **0.05034** | **-50817.587579** | **0.05147** | **0.04382** | **0.04** |
| **ND3** | **-15231.118744** | **0.04616** | **-15225.417243** | **0.04315** | **0.06974** | **<0.001** |
| ND4 | -63386.712570 | 0.03342 | -63386.654004 | 0.03330 | 0.03417 | 0.73 |
| ND4L | -13070.165371 | 0.03490 | -13069.906337 | 0.03433 | 0.03848 | 0.47 |
| **ND5** | **-85795.088154** | **0.04241** | **-85792.436300** | **0.04158** | **0.04770** | **0.02** |
| ND6 | -24185.412225 | 0.05868 | -24185.292670 | 0.05823 | 0.06181 | 0.62 |

**Supplementary Table S7.**
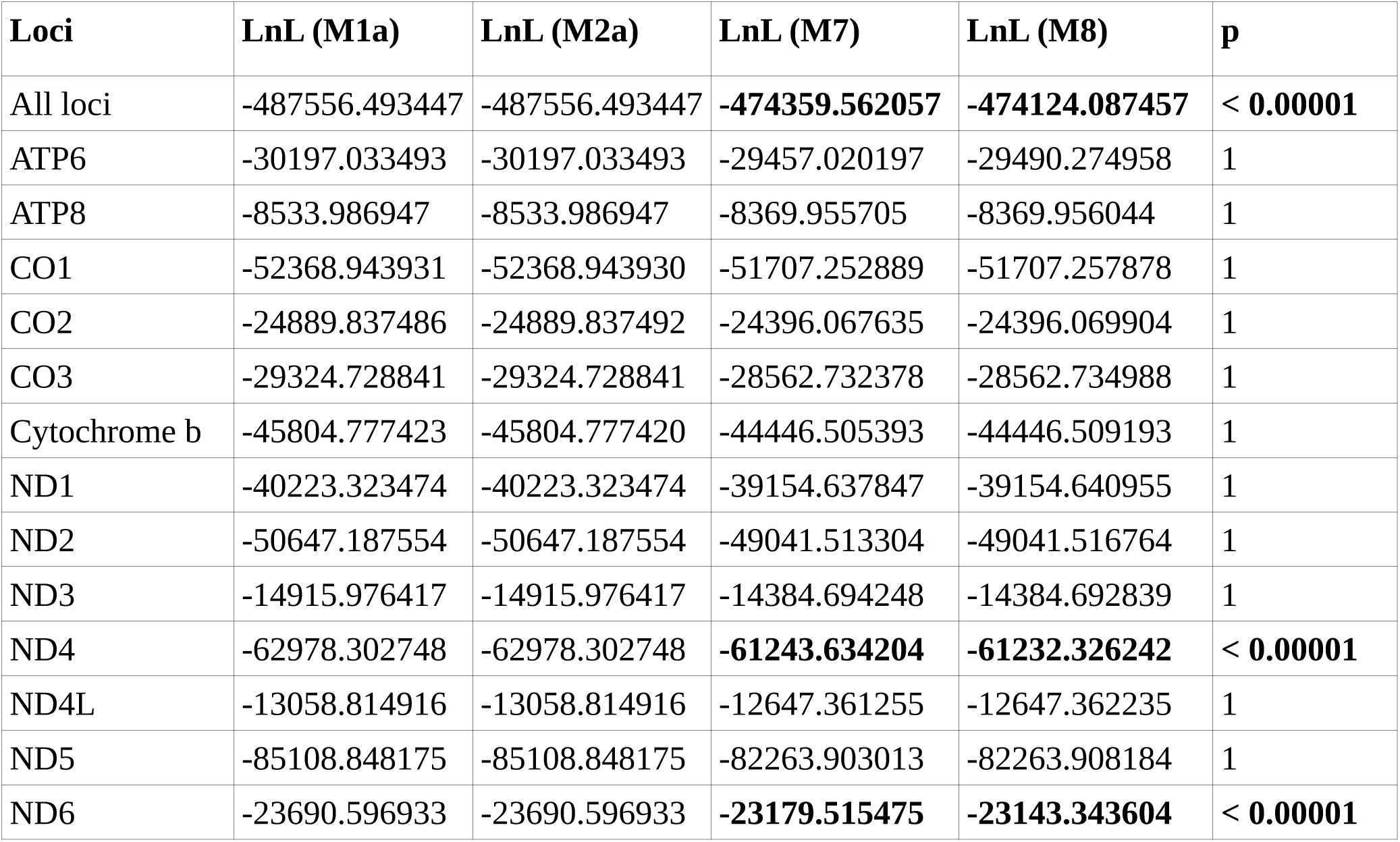
M1a/M2a and M7/M8 models comparisons to detect positive selection (site model). The all loci analyses was performed on the rooted tree. Models in bold represent comparisons where the alternative model was significantly better (at the 0.05 threshold). None of the M1a/M2a comparisons were significant (p=1 in all analyses)

**Supplementary Table S8.** Effect of the climatic regime on positive selection (branch site test of positive selection (model=2, NSsites=2). The null model was constrained using the settings fix_omega = 1 and omega =1 whereas the alternative model was specified with fix_omega=0 (that is omega is estimated)

| Loci | LnL (null model) | LnL (alternative model) | p |
| --- | --- | --- | --- |
| All loci | -487464.299047 | -487464.299058 | 1 |
| ATP6 | -30194.628880 | -30194.628880 | 1 |
| ATP8 | -8533.986949 | -8533.986947 | 1 |
| CO1 | -52368.940744 | -52368.940744 | 1 |
| CO2 | - 24883.519409 | -24883.519409 | 1 |
| CO3 | -29314.368048 | -29314.368048 | 1 |
| Cytochrome b | -45804.728830 | -45804.728831 | 1 |
| ND1 | -40218.364284 | -40218.364284 | 1 |
| ND2 | -50634.167821 | -50634.167821 | 1 |
| ND3 | -14904.875182 | -14904.875182 | 1 |
| ND4 | -62972.459483 | -62972.459482 | 1 |
| ND4L | -13057.551770 | -13057.551770 | 1 |
| ND5 | -85079.811665 | -85079.811665 | 1 |
| ND6 | -23690.594198 | -23690.594198 | 1 |

**Supplementary Table S9.** Effect of the climatic regime (alternative coding) on positive selection (branch site test of positive selection (model=2, NSsites=2). The null model (model A1) was constrained using the settings fix_omega = 1 and omega =1 whereas the alternative model (model A) was specified with fix_omega=0 (that is omega is estimated).

| **Loci** | **LnL (Null model)** | **LnL (Alternative model)** | **p** |
| --- | --- | --- | --- |
| All loci | -487474.072317 | -487474.072330 | 1 |
| ATP6 | -30194.342630 | -30194.342630 | 1 |
| ATP8 | -8533.833842 | -8533.833842 | 1 |
| CO1 | -52368.940766 | -52368.940775 | 1 |
| CO2 | -24882.594996 | -24882.594996 | 1 |
| CO3 | -29313.602721 | -29313.602721 | 1 |
| Cytochrome b | -45804.776345 | -45804.776346 | 1 |
| ND1 | -40221.442919 | -40221.442919 | 1 |
| ND2 | -50635.223005 | -50635.223005 | 1 |
| ND3 | -14903.710489 | -14903.710488 | 1 |
| ND4 | -62974.509426 | -62974.509426 | 1 |
| ND4L | -13055.088330 | -13055.088331 | 1 |
| ND5 | -85081.802226 | -85081.802227 | 1 |
| ND6 | -23690.594198 | -23690.594198 | 1 |

